# Identification of potential microRNAs associated with Herpesvirus family based on bioinformatic analysis

**DOI:** 10.1101/417782

**Authors:** Kevin Lamkiewicz, Emanuel Barth, Emanuel Barth, Bashar Ibrahim

## Abstract

MicroRNAs (miRNAs) are known key regulators of gene expression on posttranscriptional level in many organisms encoded in mammals, plants and also several viral families. To date, no homologous gene of a virus-originated miRNA is known in other organisms. To date, only a few homologous miRNA between two different viruses are known, however, no gene of a virus-originated miRNA is known in any organism of other kingdoms. This can be attributed to the fact that classical miRNA detection approaches such as homology-based predictions fail at viruses due to their highly diverse genomes and their high mutation rate.

Here, we applied the virus-derived precursor miRNA (pre-miRNA) prediction pipeline ViMiFi, which combines information about sequence conservation and machine learning-based approaches, on Human Herpesvirus 7 (HHV7) and Epstein-Barr virus (EBV). ViMiFi was able to predict 61 candidates in EBV, which has 25 known pre-miRNAs. From these 25, ViMiFi identified 20. It was further able to predict 18 candidates in the HHV7 genome, in which no miRNA had been described yet. We also studied the undescribed candidates of both viruses for potential functions and found similarities with human snRNAs and miRNAs from mammals and plants.

## 1 Introduction

MicroRNAs (or miRNAs) are small RNA molecules (~20–24 nt) involved in the regulation of gene expression by targeting messenger RNAs (mRNA) for cleavage or translational repression. Usually, two processing steps are required to generate miRNAs. First, the nuclear RNase III Drosha cuts the primary miRNA which leads to the precursor miRNA (pre-miRNA)^1–3^. The pre-miRNA is subsequently transported to the cytoplasm and further processed by Dicer, another RNase III protein, to the double-stranded miRNA/miRNA*- complex^4–6^. After separating the mature miRNA from the complementary miRNA* sequence, it is loaded into the RNA-induced silencing complex (or RISC). The loaded RISC complex targets messenger RNA (mRNA) very specifically, depending on the sequence of the miRNA. An interaction between RISC and mRNA inhibits the translation of the targeted mRNA^7^.

Aside from eukaryotes, viruses can also encode miRNAs in their genome^8,9^. Over the last decade, several hundred viral miRNAs have been discovered, most of them in DNA viruses ^10–14^. Viral genomes tend to be highly diverse (especially in RNA virus) and/or adapted to the host organism (e.g. in terms of codon usage) ^15–17^. As depicted in Figure 1, viral miRNAs are shown to not only regulate viral mRNAs but also cellular host mRNAs ^18–20^.

**Figure 1:**
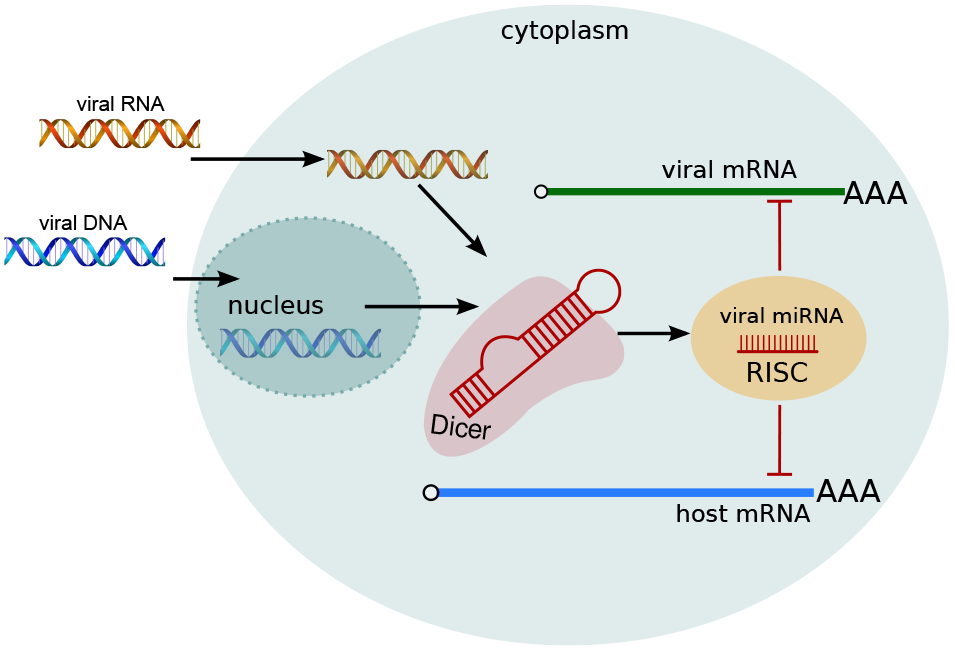
Once the viral genome entered the host cell, transcription, translation and processing mechanisms of the host are exploited. Viral precursor miRNAs are processed by Dicer and loaded into the RISC complex. Interaction with both the viral and the host mRNA is possible.

Identifying miRNAs accurately is a challenging task that requires the integration of experimental approaches with computational methods. Several miRNA prediction algorithms have been developed, but unfortunately none is able to provide a list of miRNA candidates for viruses. These prediction algorithms can be categorized into two groups. The first group deals with homology information of closely related species and compares the query sequence with annotations of known miRNAs in these species. Examples for homology-based tools are miRscan ^21^, miRSeeker ^22^ and miRAlign ^23^. The second group, including tools like TripletSVM ^24^, NOVOMIR ^25^ and HHMMiR ^26^, uses machine learning where known miRNAs are used to train a classification model and eventually to predict new candidates in query sequences.

However, as shown in Figure 2, both approaches fail for viruses (especially RNA viruses), due to the high diversity of their genomes and the lack of sufficient data.

**Figure 2:**
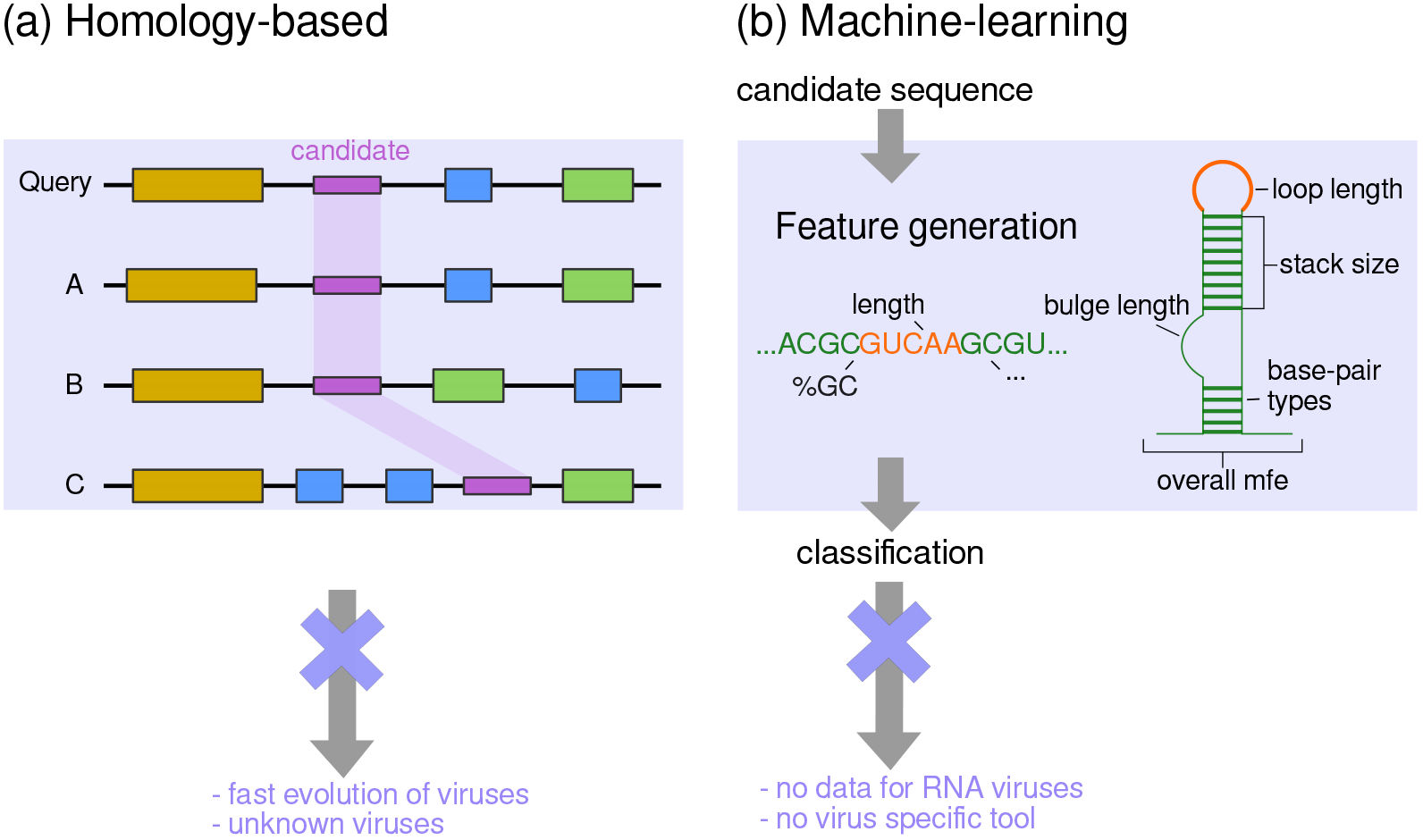
Common approaches for the prediction of miRNAs and their drawbacks on viral data. For both, the homology-based and the machine learning-based approach, the lack of data and the high diversity of viral genomes make these analyses difficult. In our comparison, homology-based approaches without the usage of synteny were studied.

Recently, we developed a pipeline for prediction of viral precursor miRNAs *de novo*, called ViMiFi, (*vi*ral *mi*RNA *fi*nder) [submitted elsewhere]. The pipeline combines homology-based and machine learning-based approaches and has been tested with six different classifier. So far, other machine learning approaches predict pre-miRNA in single sequences ^24–26^, whereas our approach processes multiple sequence alignments of viral genomes, enabling a larger feature space for classification.

In this study, we present the results of the first application of ViMiFi, applied on Human Herpesvirus 7 (HHV7). This virus has a double-stranded DNA (dsDNA) genome with 145 kb in size and belongs to the genus of *Roseoloviruses* and the (sub-)family of *Herpesviridae*. There are nine different herpesviruses known that infect human. To date, most of these nine viruses encode for viral miRNAs ^8,9,11,27–30^. The Varicella-Zoster virus encodes several small ncRNAs ^29^, however, no miRNA was cloned yet. For the HHV7 no miRNAs have been reported either.

HHV7 infections are associated with a number of symptoms such as acute febrile respiratory disease, fever, diarrhea and low lymphocyte counts ^31^. Furthermore, there are indications that HHV7 could contribute to the development of drug-induced hypersensitivity syndrome ^32^, encephalopathy ^33^, hemiconvulsion-hemiplegia-epilepsy syndrome ^34^ and hepatitis infection ^35^. Thus, HHV7 has a high relevance and identifying viral miRNAs within the HHV7 genome may lead to new therapies against it. For other human herpesviruses it has been shown that viral miRNAs may play an important role in maintaining the latency phase during virus infection ^36^.

Here, we identified 18 new regions of the HHV7 genome containing potential pre-miRNAs and searched for homologous sequences in other species from all kingdoms. In order to measure the quality of our predictions, we scanned the Epstein-Barr virus (EBV). This virus is known to encode for at least 25 pre-miRNAs. From these, ViMiFi identified 20 pre-miRNAs and further proposes 41 novel candidates within the EBV genome. Figure 6 gives a schematic overview of the genome organization of EBV with the positions of our predicted candidates.

## 2 Methods

### Virus Sequence Data

We used eleven different HHV7 genome sequences for our analyses. All sequences were downloaded from the ViPR database ^37^ (see Table 1). The multiple sequence alignment (MSA) was created with mafft^38^, containing 163.407 columns and an average number of 259 gaps per sequence.

**Table 1:** Overview of HHV7 genomes used in this study. Sequences were downloaded from the ViPR database ^37^.

| Accession ID | Sequence length |
| --- | --- |
| NC_001716 | 153.080 |
| AF037218 | 153.080 |
| DD138892 | 153.080 |
| DJ052727 | 153.080 |
| DL242081 | 153.080 |
| FV537034 | 153.080 |
| FV537035 | 153.080 |
| U43400 | 144.861 |
| DD250331 | 144.861 |
| DI115158 | 144.861 |
| HV987120 | 144.861 |

### Training data

Our positive training data consists of all viral precursor miRNAs (320) that are publicly available in miRBase 22.0^39^. For each viral precursor structure of the positive set we randomly sampled three sequences from the whole genome of the same virus. The length of such a sequence was randomly set between 70 and 120 nucleotides. The sampling procedure was repeated until three negative instances per precursor were found. Each sampled sequence had to fold into a characteristic stem-loop structure of minimum size of 50 nucleotides and at least 14 base pairings. Furthermore, the minimum free energy (MFE) of the sampled sequences had to be lower than 18 kcal/mol. These parameters were chosen as they represent a canonical pre-miRNA. Performing an RNAclust ^40^ analysis on clustered viral pre-miRNAs indicates that the consensus structure of 157 pre-miRNA sequences fulfill the described characteristics. The structure is shown in Figure 3. As a final filter, we checked whether the sampled sequence had a sequence similarity over 90 % with one of the known precursor sequences. Sequence similarity was calculated with the Levenshtein distance ^41^.

**Figure 3:**
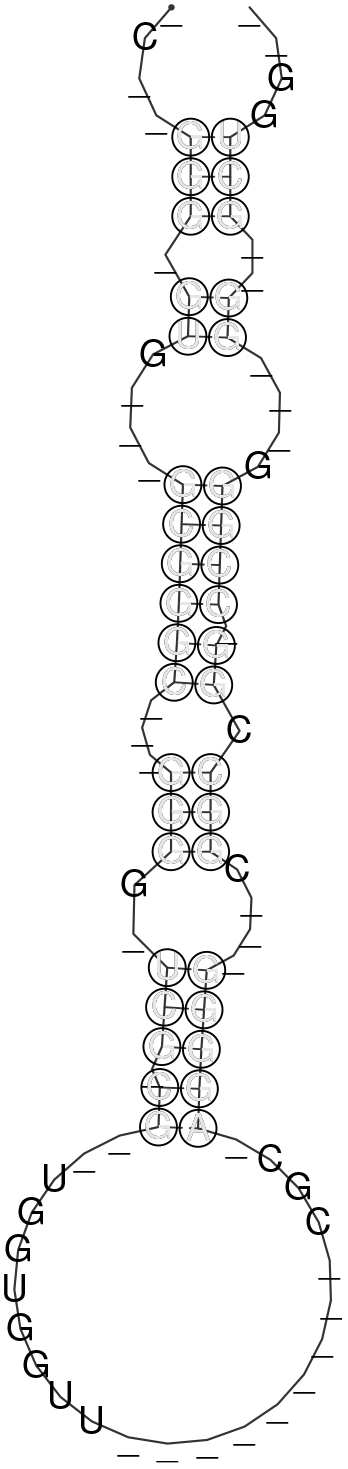
Consensus structure of 157 viral pre-miRNAs clustered with RNAclust and folded with RNAalifold. Due to the features of this structure we set the sampling parameters for negative instances as described.

### Feature discrepancy

Properties derived from the sequence and the structure of a query RNA are called features. Here, we used triplet frequencies (introduced in ^24^), relative GC-content of the sequence, the MFE of the structure, number and length of bulges, the number of pairings in the largest stacking and the loop length of the structure. In order to analyze which feature is important for the classification we performed a F-value calculation (see Equation 1), where *x̅_i_*^(+)^, *x̅_i_*^(*−*)^ and *x̅_i_* are the averages of the *i*th feature of the positive, negative and the whole dataset, respectively. Furthermore, *n*_+_ and *n_−_* denote the number of instances in the positive and negative set.

### Classification by ViMiFi

Here, we used the alignment mode of ViMiFi v.0.1. MSAs have to be provided to ViMiFi. As ViMiFi is not published yet, we briefly describe the workflow and default parameters used in this study. The MSA is processed with a sliding window of size 120 and stepsize 20. Each window is folded with RNAalifold 2.4.9 ^42^ from the ViennaRNA package.

We use the combination of mafft and RNAalifold since we are looking in complete viral genome alignments for pre-miRNAs. Algorithms like LocARNA ^43^ do consider sequence and structural information for the creation of an alignment simultanously, however, it is not feasible to create such an alignment for several genomes of over 150.000 nucleotides in size.

Each window of our MSA is then shuffled column-wise 1.000 times. The shuffled window is folded in the same way as the original MSA fragment. From the resulting structures a p-value is calculated based on the z-score of the MFE structures.

Among the features mentioned above, ViMiFi relies on so-called triplet features, proposed in TripletSVM 24. Triplets are suitable to model the sequence and local structure of a query sequence. For each triplet of nucleotides, the second nucleotide itself and the information whether a nucleotide of a triplet is paired or unpaired in the predicted secondary structure is stored. Thus for example, the sequence GGGGUCCCC with the structure ((((…)))) in dot-bracket format would lead to the following triplets: 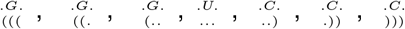.

Parameters of ViMiFi were set to their respective default values (windowsize: 120, stepsize: 20, number of shuffled sequences: 1.000), with the exception of the p-value cutoff, which was set to *p <* 0.01.

### Secondary Structure analysis of single candidates

Potential candidates for viral pre-miRNAs were analyzed regarding their secondary structure. For this, we applied RNAfold 2.4.9^42^ with parameters –noLP -p on each candidate sequence and colored the resulting MFE structure based on the base pairing probabilities observed in the Boltzmann distribution of secondary structures.

### Homolog Search & Function

An exhaustive search on the Rfam database version 13.0^44,45^ was performed using Infernal 1.1.2^46^ to annotate some of the predicted candidates functionally. Using cmscan from the Infernal package with default parameters, all potential pre-miRNA of HHV7 and EBV were compared with the covariance models of the Rfam release. Furthermore, for each pre-miRNA candidate a BLAST 2.7.1+^47^ search was performed against all miRBase precursor sequences. For this, we reduced the word size to 12 for the blast search, thus resulting in the command blastn –word_size 12.

### Validation of predictions using small RNA-Seq data

In order to see whether candidates predicted in the HHV7 by ViMiFi are already supported by publicly available RNA-Seq data, we used the RNA-Seq libraries of the recent publication by Lewandowska *et al.* (2017) ^48^. The libraries are accessible via http://doi.org/10.5281/zenodo.400950. Viral reads were mapped with TopHat2 v2.1.1^49^ with default parameters against the HHV7 reference genome (NCBI accession NC_001716) and data was displayed using the Integrative Genome Browser (IGV) ^50^.

## 3 Results

### Known EBV pre-miRNAs are predicted by

ViMiFi We examined several different Epstein-Barr virus genomes to identify pre-miRNAs. For this, all EBV-derived pre-miRNAs were excluded

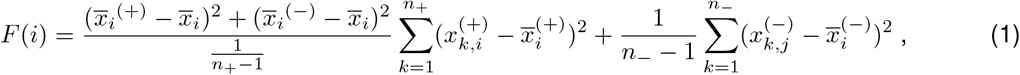

from our training set to avoid an overfitting effect with regard to EBV. Figure 6 gives an overview of the coding sequences (CDS) of both strands within the EBV genome, the 25 known pre-miRNAs annotated in miRBase and all 61 candidates predicted by ViMiFi.

We have successfully predicted 20 of 25 known pre-miRNAs curated in miRBase. The five missing pre-miRNAs contain three features being different from the other 315 of the positive set pre-miRNAs: the triplet 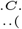 the triplet 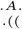 and the loop length. Figure 4 visualizes these differences based on our F-value analysis. On average, the loop length of missed pre-miRNAs is about 6 nucleotides smaller, whereas the frequencies of the two triplets are higher compared to the identified pre-miRNAs. We hypothesize that ViMiFi cannot identify these five missing pre-miRNAs, because their features are not similar enough to the ones learned from our positive training set. Reevaluating the feature design and feature selection of ViMiFi, as well as obtaining more validated viral pre-miRNAs for the training set, may lead to even better predictions, and thus increases the precision.

**Figure 4:**
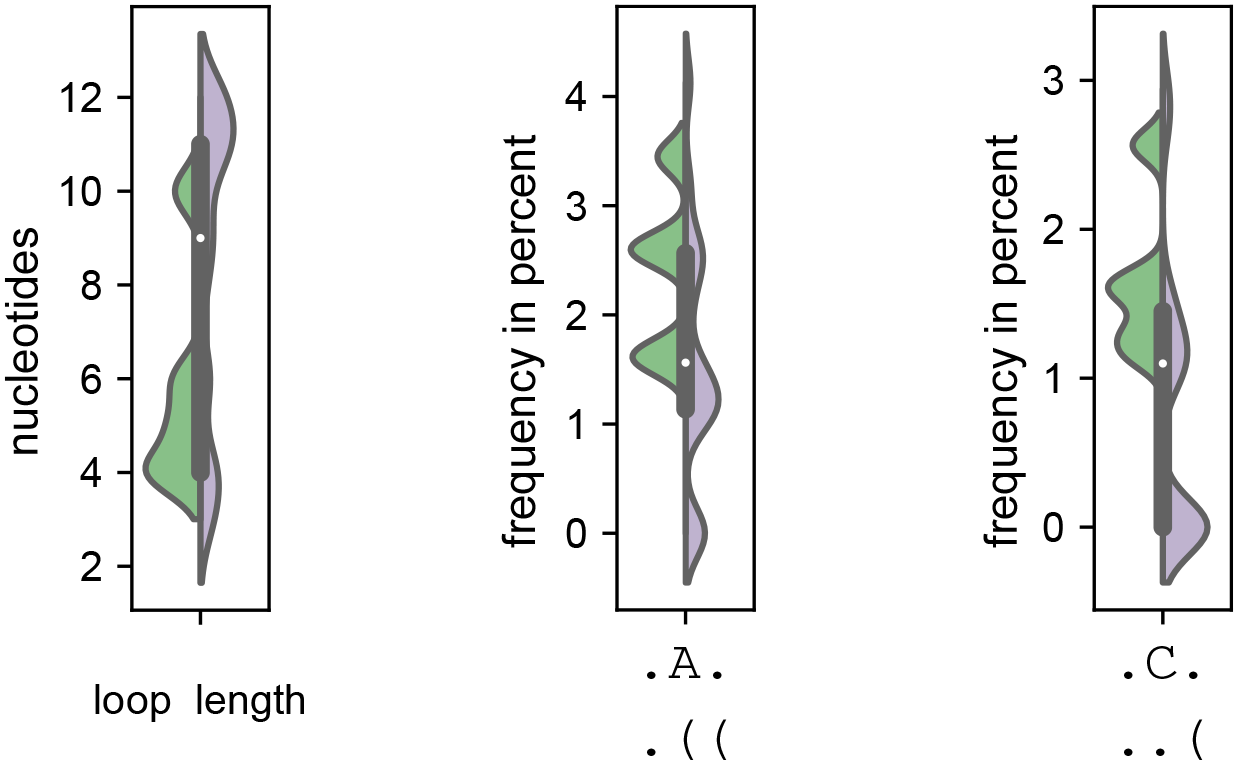
Violinplots of the three features with the highest F-value between missing (green) and identified (purple) EBV pre-miRNAs. Missed pre-miRNAs tend to have a smaller loop region as well as higher frequencies of the two triplets 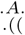 and 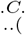.

We compared the start and stop positions of our predicted EBV candidates with the genomic coordinates of annotated pre-miRNAs of miRBase. This comparison revealed that 20 known pre-miRNAs were identified by ViMiFi.

### ViMiFi predicted 41 novel pre-miRNA candidates

We predicted with ViMiFi 41 novel pre-miRNA candidates for the EBV genome and investigated them further with Infernal and blastn. As shown in Figure 6, the majority of our predictions cluster in three regions. These regions do not have a CDS on either strand in the NCBI annotation and two of these regions are already known to be miRNA clusters – namely the BHRF1 and the BART cluster. However, one cluster of our predictions (around position 7.800 – 8.100) neither aligns with known pre-miRNAs nor with CDS regions. The candidates miRNA_ebv_03 to miRNA_ebv_07 belong to this cluster. None of these candidates have a homolog pre-miRNA in the miRBase. Thus, we extracted the region from position 7.800 – 8.100 of the EBV genome and made a structure analysis using RNAfold ^42^. The MFE structure is shown in Figure 5 and resembles the shape of a potential primary miRNA (pri-miRNA) ^51–53^. For all results of the blastn search we refer to Supplement Table S1.

**Figure 5:**
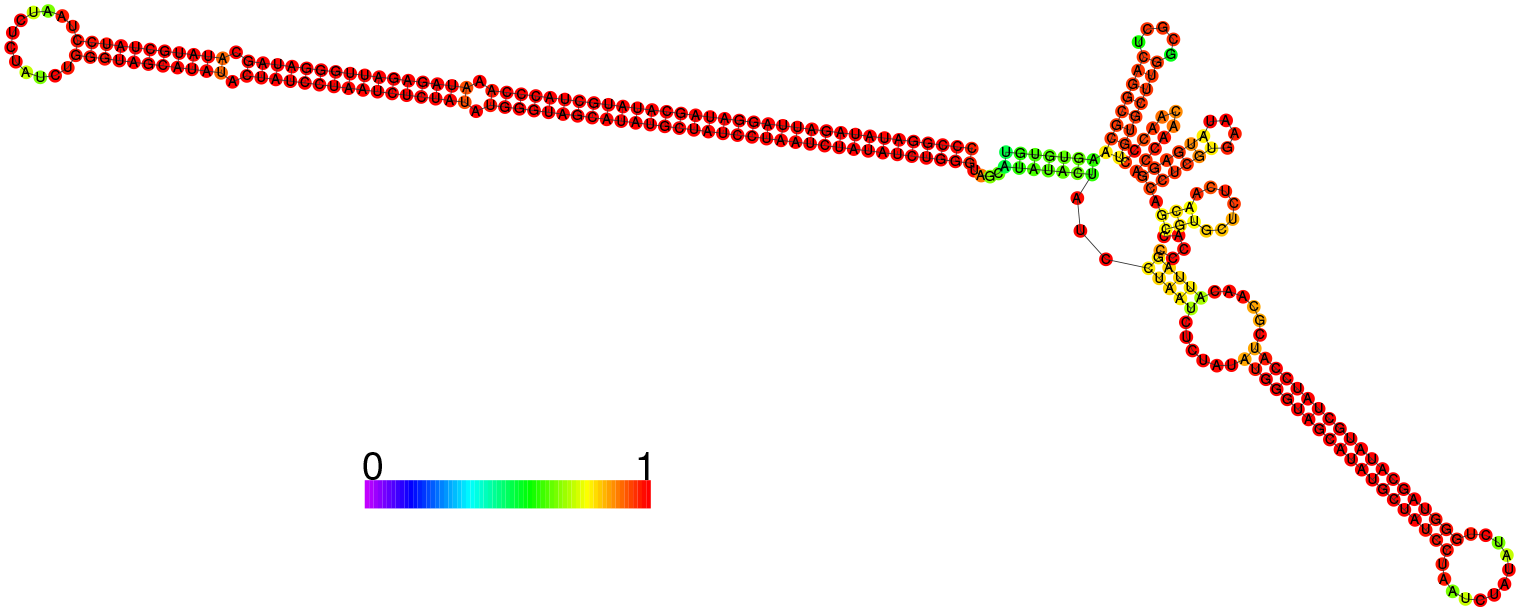
MFE structure of the region 7.800 – 8.100 of the EBV genome, folded with RNAfold and colored based on the base pairing probabilities observed in the set of sampled structures in the Boltzmann ensemble. The structure resembles a plausible pri-miRNA.

**Figure 6:**
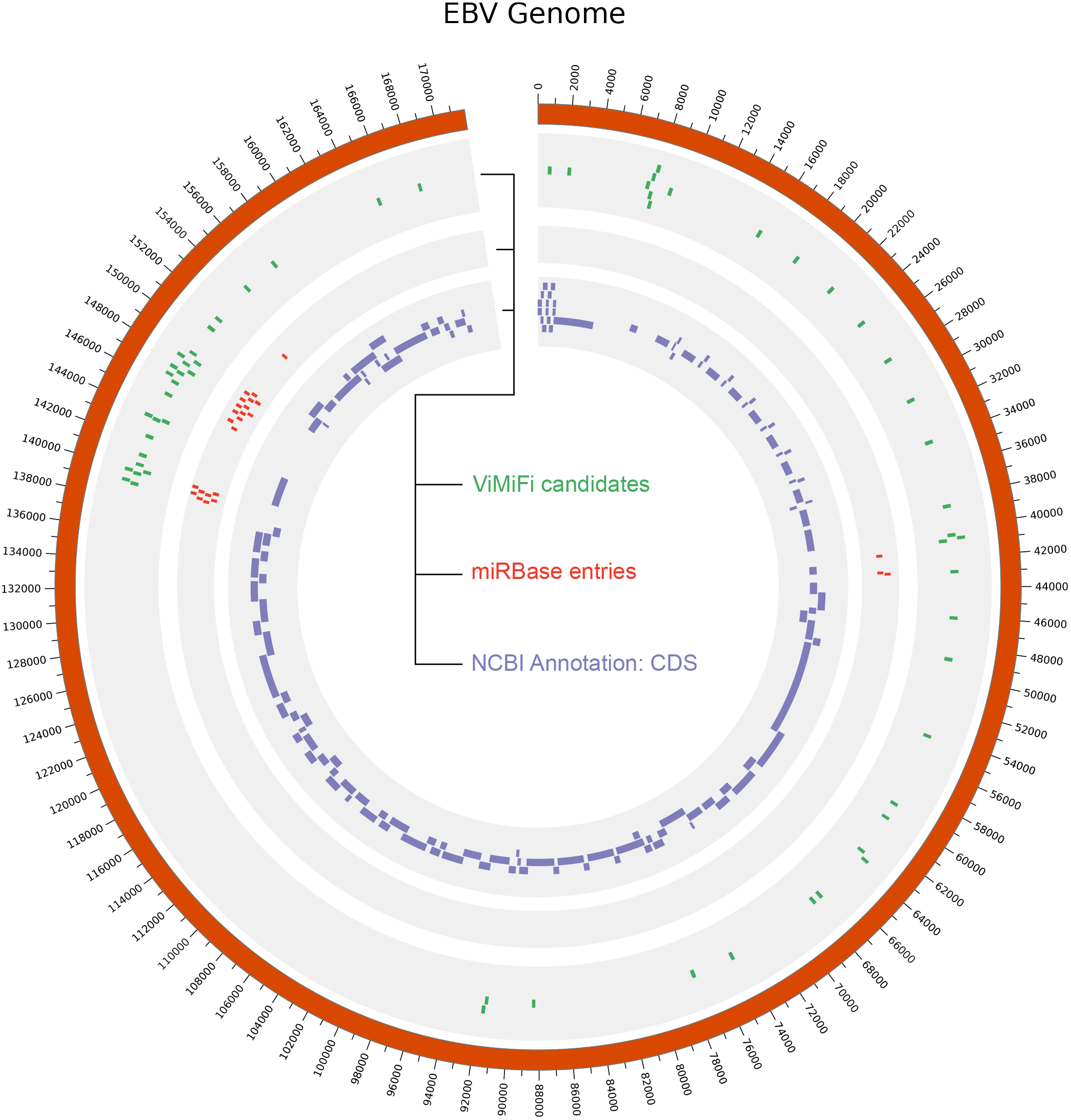
Circular representation of the Epstein-Barr virus genome and annotation of known and predicted pre-miRNAs. The inner track in purple represent annotated coding regions from NCBI. The middle track shows all pre-miRNA entries of the miRBase in red. All candidates predicted by ViMiFi are represented in green in the outer track.

Applying a covariance model search from the Infernal toolkit against all known RNA families of the Rfam database revealed 37 hits with a significant (*<* 0.01) e-value. The results of this search are shown in Table 2. Interestingly, the candidates miRNA_ebv_03 to miRNA_ebv_07 are not associated with EBV. However, all of them have similarities with the RNA family miR-563. This family is conserved among mammals, however, the precise function has not been identified, yet. Since the e-values of these hits are close to 1, the predicted EBV pre-miRNA cluster does probably not belong to this RNA family, but might resemble its own RNA family.

**Table 2:**
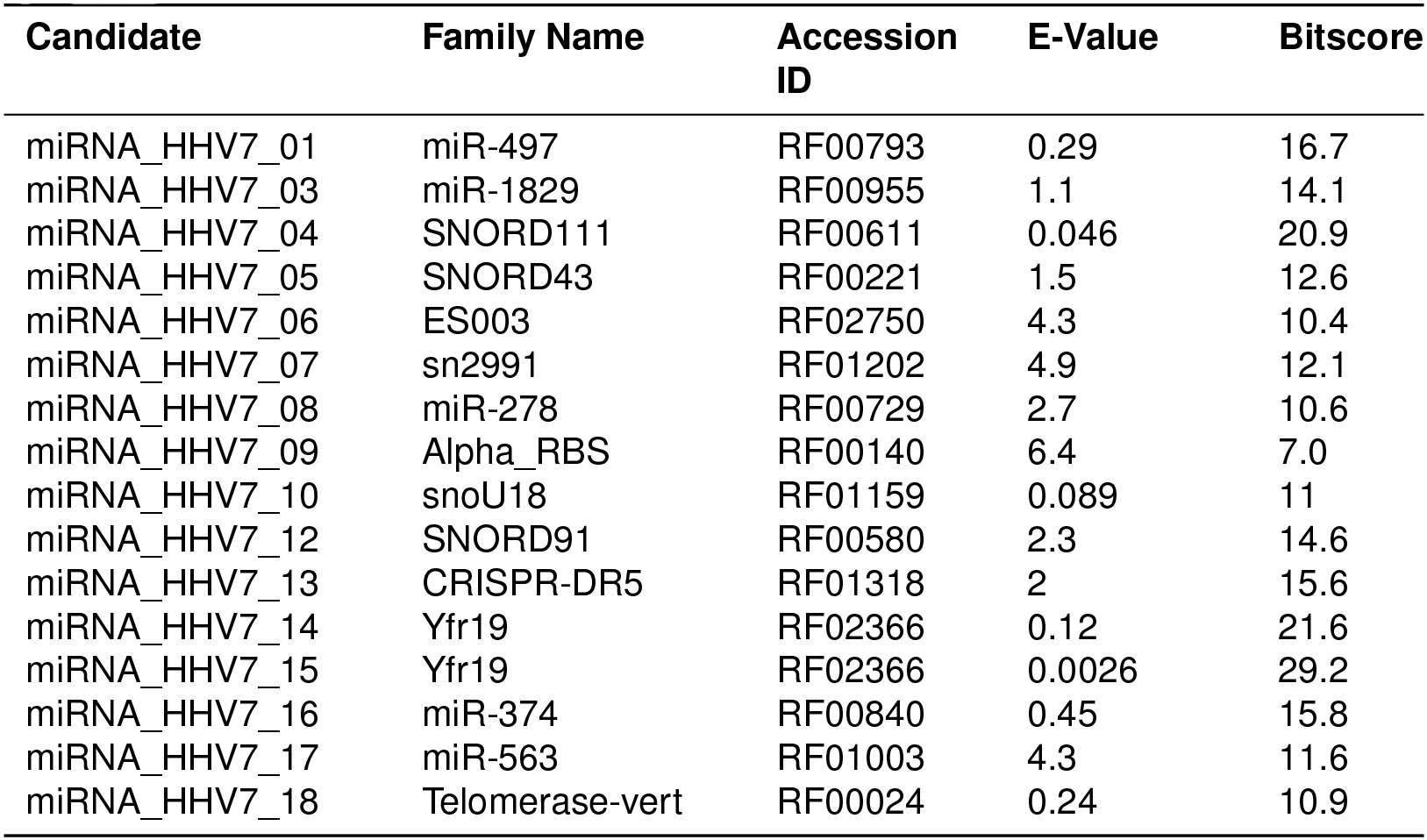
Results of the covariance model search with Infernal applied to predicted EBV candidates. We searched against the Rfam database. For the complete list of all Infernal results we refer to Supplement Table S2.

Furthermore, the candidates miRNA_ebv_09 to miRNA_ebv_15 have hits on the RNA family ebv-sisRNA-2, which is not a miRNA, but a long non-coding RNA (lncRNA) of EBV. ViMiFi detected these six candidates in the region of ebv-sisRNA-2, because this lncRNA has many stable local stem-loops in its secondary structure which are similar to pre-miRNAs. The ebv-sisRNA-2 lncRNA may play an important role in the maintenance of virus latency ^54,55^.

### 18 novel pre-miRNAs in HHV7

`ViMiFi identified` 18 different regions in the HHV7 alignment, consisting of the genomes as displayed in Table 1, to be potential pre-miRNAs. Table 3 shows all candidates sorted by their genomic coordinates relative to the reference genome and Figure 7 shows the positions of the candidates in relation to the CDS annotation of NCBI.

**Table 3:** Overview of predicted pre-miRNAs in HHV7 using ViMiFi. Each candidate is assigned an unique ID. The start and stop positions are relative to the reference genome accessible via the NCBI accession NC_001716.

| Candidate | Start Position | Stop Position |
| --- | --- | --- |
| miRNA-HHV7_01 | 9.663 | 9.761 |
| miRNA-HHV7_02 | 28.215 | 28.316 |
| miRNA-HHV7_03 | 34.374 | 34.467 |
| miRNA-HHV7_04 | 51.590 | 51.668 |
| miRNA-HHV7_05 | 61.730 | 61.807 |
| miRNA-HHV7_06 | 67.481 | 67.585 |
| miRNA-HHV7_07 | 75.954 | 76.024 |
| miRNA-HHV7_08 | 87.892 | 87.995 |
| miRNA-HHV7_09 | 94.797 | 94.881 |
| miRNA-HHV7_10 | 104.503 | 104.588 |
| miRNA-HHV7_11 | 110.869 | 110.943 |
| miRNA-HHV7_12 | 116.193 | 116.306 |
| miRNA-HHV7_13 | 122.944 | 123.026 |
| miRNA-HHV7_14 | 127.429 | 127.537 |
| miRNA-HHV7_15 | 127.487 | 127.595 |
| miRNA-HHV7_16 | 127.723 | 127.813 |
| miRNA-HHV7_17 | 148.530 | 148.625 |
| miRNA-HHV7_18 | 152.705 | 152.811 |

**Figure 7:**
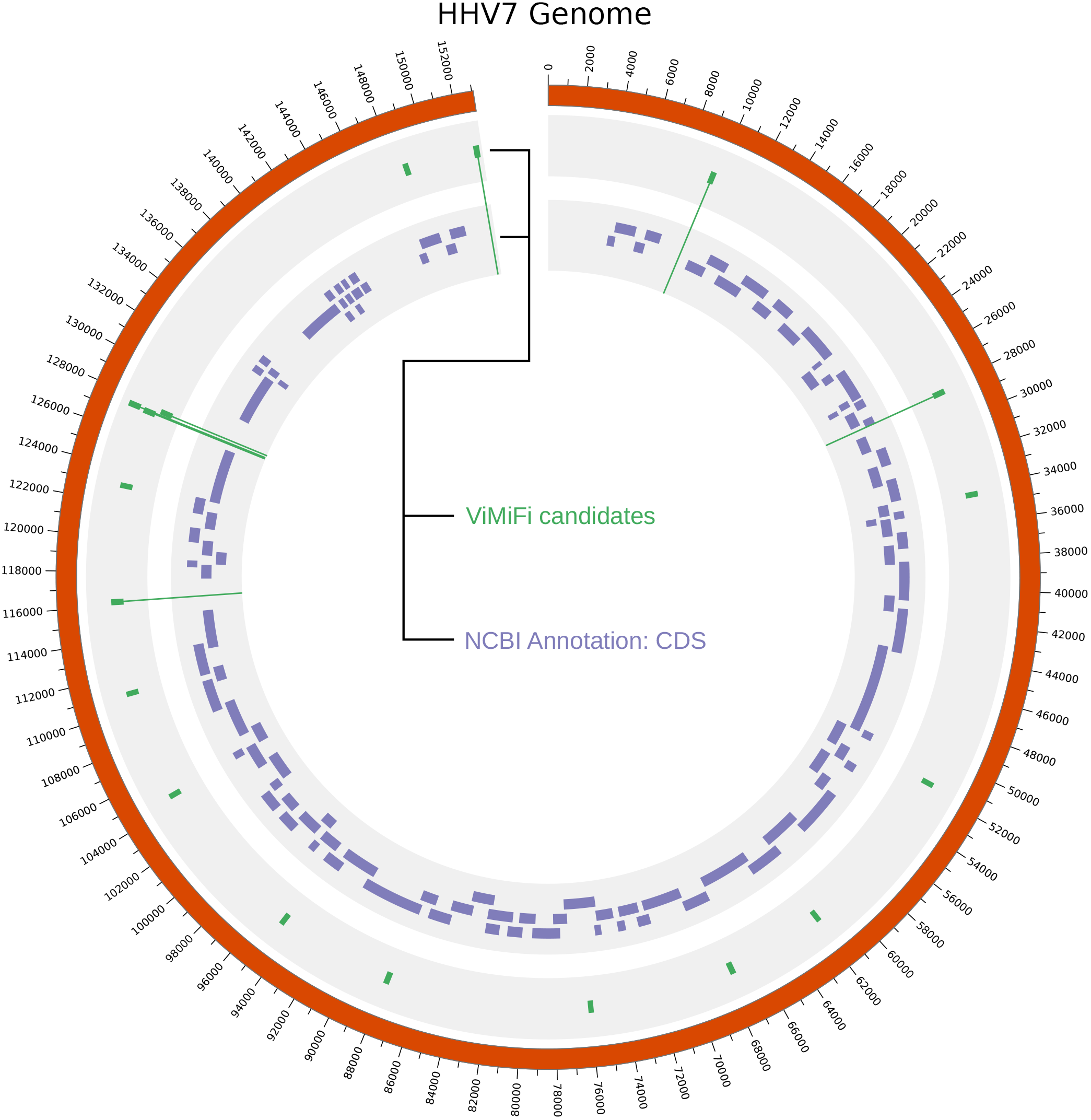
Circular representation of the Human Herpesvirus 7 genome and annotation of predicted pre-miRNAs. The inner track in purple represent annotated coding regions from NCBI. All candidates predicted by ViMiFi are represented in green in the outer track. Lines from candidates going into the inner track indicate potential pre-miRNAs that do not overlap with annotated CDS regions.

Each candidate was analyzed with `blastn` and `Infernal`. Performing the blastn search against the `miRBase` database led to some hits in several plants, *Homo sapiens* and *Macaca mulatta* (see Supplement Table S3), however the e-values indicate that those hits are probably not real homologs, but at least share some similarities in terms of sequence.

### Seven HHV7 candidates are not overlapping with coding regions

From our 18 predicted candidates within the HHV7 genome, 7 do not overlap with annotated CDS on any strain. These 7 candidates are miRNA_HHV7_01, miRNA_HHV7_02, miRNA_HHV7_12, miRNA_HHV7_14, miRNA_HHV7_15, miRNA_HHV7_16 and miRNA_HHV7_18. In Figure 7 the corresponding green tiles are drawn with a line into the CDS track. Even more intriguing, for the candidates miRNA_HHV7_01 and miRNA_HHV7_18, no blastn hits against the miRBase were obtained at all. Thus, no other known pre-miRNA derived from all organisms in the miRBase are similar to these new candidates. Analyzing the secondary structures of the 7 candidates shows that all of them are able to fold into a structure with at least one stable stem-loop (see Figure 8). Even structures that do not form the canonical single hairpin structure of pre-miRNAs, may exploit another pathway of pre-miRNA processing, for example the tRNase Z-dependent pathway ^56^. As first step of validation, we examined the recently published RNA-Seq data of Lewandowska *et al.* ^48^. We observed weak expression for the candidates miRNA_HHV7_02, miRNA_HHV7_12, miRNA_HHV7_14, miRNA_HHV7_15 and miRNA_HHV7_17 (see Supplement Figures S1 – S4). These observations indicate transcription activity at the predicted pre-miRNA regions, however, they are no clear evidence for the presence of pre-miRNAs. We therefore argue that these novel candidates are worth investigating by experiments in order to validate their presence and potential targets during infection of human cells.

**Figure 8:**
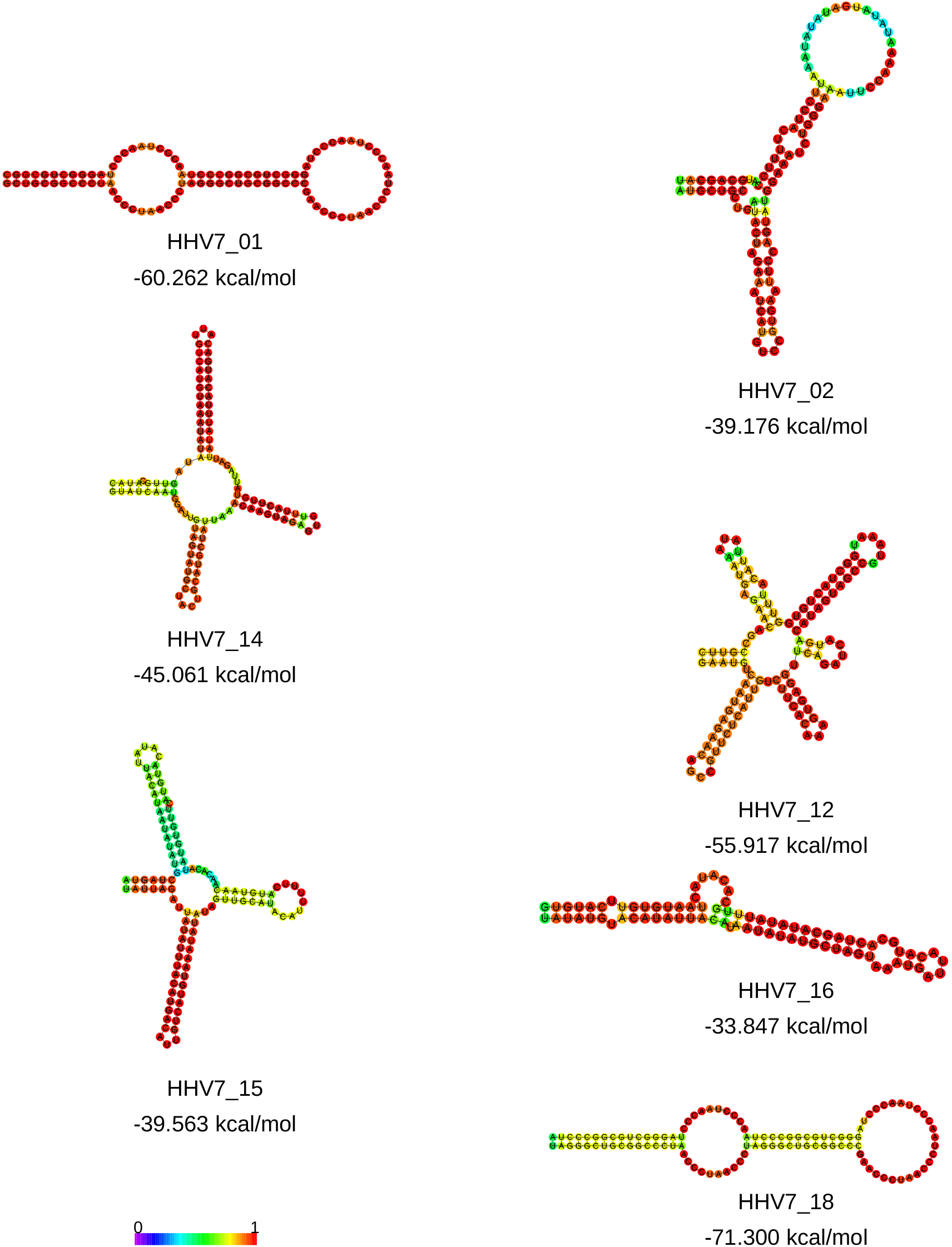
Predicted secondary structures of the potential novel pre-miRNAs encoded by HHV7. Displayed are the minimum free energy structures of all candidates that do not overlap with annotated CDS. Each structure is colored based on the base pairing probabilities derived from the partition function. The more a nucleotide is colored red, the higher the likelihood of the predicted structure. Since the candidates 02, 12, 14 and 15 do not fold like known pre-miRNAs they arguably could be false positives. However, Supplement Figures S1 – S4 show that at least some reads can be aligned to the positions of those candidates, indicating active transcription of these regions. It might be possible that those four predicted candidates are processed in alternative miRNA pathways in contrast to the canonical Drosha and Dicer pathway.

Interestingly, one of the candidates, that overlaps with CDS, miRNA_hhv7_06 has a significant blastn hit on the human miRNA hsa-miR-4432. However, this human miRNA is not confidently reported yet and has around 100 of predicted potential targets within the human transcriptome ^57^.

### Two HHV7 candidates can be linked to known snRNAs

Further, we searched for similarities of the predicted HHV7 candidates with other RNA families in the Rfam database. A co-variance model search for each candidate was performed. The most significant results are shown in Table 4, whereas all results are shown in Supplement Table S4. Only one hit achieved an e-value *<* 0.001, namely the non-overlapping miRNA candidate miRNA_hhv7_15 on the RNA family C**y**anobacterial **f**unctional **R**NA 19 (Yfr19). Yfrs are known to regulate gene expression in cyanobacteria. In particular, Yfr19 is controlled by phage infection ^58^, linking this bacterial ncRNA to virus infections.

**Table 4:**
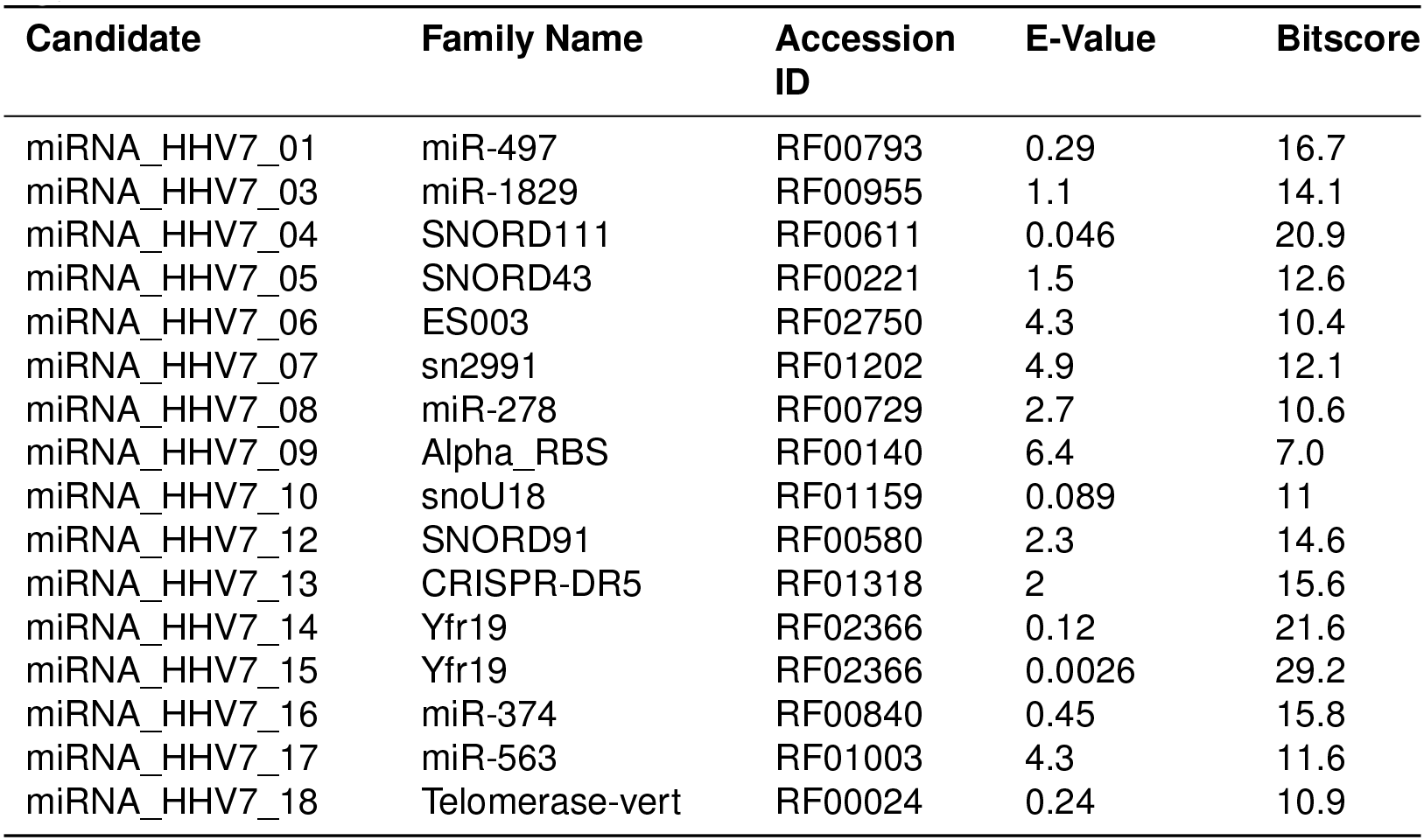
Results of the covariance model search with Infernal applied to predicted HHV7 candidates. We searched against the Rfam database. For each candidate the most significant hit is reported. For all hits we refer to Supplement Table S4.

Furthermore, miRNA_HHV7_04 has similarities to the RNA family SNORD111. SNORD RNAs are small non-coding RNAs which are involved in the modification of small nuclear RNAs (snRNAs). The SNORD111 (also known as HBII-82) is being predicted to guide the 2’O-ribose methylation of 28S rRNA in mouse, human, other mammals and aves 59,60. The Infernal search also found similarities between the RNA family snoU18, which is mainly observed in insects, and the candidate miRNA_HHV7_10. The snoU18 is known to be a C/D-box snRNA and is associated with methylation as well. The predicted candidates may compete with these cellular snRNAs to prevent methylation. Moreover, there exists a non-canonical pathway for pre-miRNA processing that is derived from C/D-box snRNAs ^56^.

## 4 Conclusion

We applied the virus specific pre-miRNA prediction pipeline ViMiFi on the human herpesvirus 7, the solely human-infecting herpesvirus that has no miRNAs described, and on Epstein-Barr virus, a human-infecting herpesvirus with 25 described pre-miRNAs. We were able to identify 20 out of these 25 precursor structures, and further, predict 41 more regions to be potential pre-miRNAs candidates. Out of these 41 candidates, 5 do not overlap with any known annotation, neither the NCBI coding region nor the miRBase entries.

Moreover, we proposed 18 pre-miRNAs candidates in HHV7. Out of these, 16 candidates show similarities to known pre-miRNAs in plants, *Homo sapiens* and *Macaca mulatta*, whereas the candidates miRNA_HHV7_01 and miRNA_HHV7_18 did not yield any result in the applied homology search. Further, seven candidates do not overlap with known annotated protein coding genes. The predicted secondary structures and the genomic context of these seven candidates indicate that they are undescribed pre-miRNAs encoded by Human Herpesvirus 7. Even though, some structures do not resemble the canonical pre-miRNA, they could be processed via non-canonical pathways like the tRNase-Z or snRNA-derived pathway.

Comparisons with RNA families stored at the Rfam database show minor similarities with other miRNA and snoRNA families. These findings suggest that the candidates predicted by ViMiFi may have similar functions. Intriguingly, two HHV7 candidates have similarities to C/D-box snRNAs, which are known to guide the methylation of rRNA. These methylation may lead to “specialized ribosomes” ^61^. We hypothesize that viral ncRNAs with similarities to such cellular snRNAs may compete with these snRNAs and prevent the creation of specialized ribosomes, and thus, ensure viral protein translation.

The next obvious step is the validation of our candidates *in vitro* and/or *in vivo*. Since the data of viral pre-miRNAs is limited, and thus the performance of ViMiFi is limited as well, we hope that virologists are intrigued by these findings and are open for collaborations – each validated pre-miRNA can refine the model used by ViMiFi and thus lead to more accurate results in the future.

## Author Contributions

KL, BI and MM conceived of the presented idea. KL performed the computations. KL, EB and BI analyzed the data and discussed the results. EB performed the computational RNA-Seq validations. KL and BI wrote the manuscript with critical input from EB and MM.

## Acknowledgments

The work of KL is funded by the German Federal Ministry for Higher Education and Research (BMBF): STIKO Serologie (InfectControl 2020), Project Number 03ZZ0820A. We thank Anna Strototte and Katja Meyer for proofreading the manuscript.

